# Cryptic variation alters gene dosage sensitivity to shape inflorescence architecture in tomato

**DOI:** 10.64898/2026.05.07.722400

**Authors:** Gwen Swinnen, Stéphanie Afonso, Elia Lacchini, Stéphanie Stolz, Eléonore Lizé, Sebastian Soyk

## Abstract

Phenotypic diversity arises in large part from genetic variants at multiple interacting loci, many of which alter gene dosage rather than abolish gene function. Dosage-sensitive variants, which often produce nonlinear phenotypic outcomes, can be exploited to fine-tune quantitative traits for crop improvement using genome editing. However, the phenotypic outcomes of individual variants can differ substantially across genetic backgrounds, as segregating alleles may modulate allelic effects in unexpected ways. Yet, how genetic background shapes gene dosage effects remains underexplored. Here, we show that MADS-box gene dosage effects, which can be used to tune tomato inflorescence architecture for optimal fruit yield, differ profoundly between distinct genetic backgrounds. We mapped the genetic basis of this background dependency and identified the cryptic modifier locus *suppressor of branching 2* (*sb2*), which contains the conserved floral identity gene *ANANTHA*. We show that natural variation at *sb2* modulates how inflorescence architecture responds to MADS-box dosage effects from natural and engineered loss-of-function mutations. Our findings illustrate how cryptic genetic variants can reshape gene dosage relationships and underscore the importance of characterizing such hidden variation for predictive engineering of quantitative traits using genome editing.

**Significance Statement:** Advances in crop genome editing enable precise modifications of gene dosage to fine-tune quantitative traits in crop improvement, but the predictability of such strategies remains limited. We show that hidden genetic differences, known as cryptic variation, can alter how gene dosage changes influence plant growth and development. Using tomato inflorescence architecture as a model, we characterize a natural cryptic modifier locus, *suppressor of branching 2* (*sb2*), that modifies the effects of natural and engineered mutations in dosage-sensitive MADS-box genes. Our findings demonstrate that gene dosage effects depend on genetic background and highlight an often-unrecognized constraint on precision breeding by genome editing. Accounting for similar cases of cryptic variation will be essential for predictable engineering of quantitative traits in crops.

## Introduction

A major challenge in genetics is to understand how complex genotypes shape quantitative traits. The prediction of phenotypes from genotypes is complicated by the interaction between genes, in which variation at one gene influences how another gene affects a phenotype (1–3). Genetic interactions become especially difficult to predict when they involve cryptic variants, which are alleles with little or no phenotypic effects unless revealed in distinct genetic backgrounds through interaction with other variants (4, 5). Cryptic variants can accumulate neutrally and have the potential to fuel adaptations in both natural (5) and domesticated (6–8) contexts. However, empirical evidence showing how cryptic variants involved in genetic interactions can modify quantitative traits is limited in crops.

Genetic interactions frequently occur among genes encoding transcriptional regulators or signaling pathway components, many of which form multimeric protein complexes and are sensitive to gene dosage. Genomic changes that modify gene dosage, such as duplications, deletions, polyploidization, and loss-of-function mutations, often modulate the level of functional gene product. For dosage-sensitive genes, even modest changes in protein abundance can have phenotypic effects due to nonlinear relationships between gene expression and phenotypic outcomes (9). A molecular framework for how such genes contribute to quantitative trait variation is provided by the gene balance hypothesis (10–12). Imbalances in gene dosage perturb stoichiometric relationships among components of multimeric protein complexes, thereby altering protein complex activity and ultimately downstream network dynamics. Such gene dosage effects can explain heterosis (hybrid vigor) in crops, where heterozygosity at single or multiple loci shifts network dynamics toward new optima and enhances crop performance (13–15). An illustrative example involves heterozygosity at florigen pathway genes, producing heterotic effects for flowering time and yield in tomato (*Solanum lycopersicum*) and rice (*Oryza sativa*) (16–18).

Crop breeding can leverage dosage-sensitive genetic interactions for yield improvement. In fruit crops, productivity depends on inflorescence architecture, the branching pattern of the flower-bearing shoots, which is established by the rate at which shoot apical meristems mature to floral termination (19, 20). In tomato, meristem maturation and inflorescence branching are regulated by *JOINTLESS2* (*J2*) and *ENHANCER OF J2* (*EJ2*), members of a family of MADS-box transcription factors that form tetrameric protein complexes (21–23). Tomato breeders selected a *j2* null allele (*j2^TE^*) that eliminates the fruit abscission zone and facilitates mechanical harvesting, yet does not alter inflorescence branching (24). When breeders accidentally recombined the *j2^TE^* allele with a cryptic *ej2* partial loss-of-function allele (*ej2^W^*), which produces only ∼30% of functional *EJ2* transcript, this led to excessively branched inflorescences with poor fruit set in the *j2^TE^ ej2^W^* double mutant (23). Interestingly, CRISPR-engineered *j2^CR^ ej2^CR^* null mutants branch even more strongly than the natural *j2^TE^ ej2^W^* mutant, whereas *j2^CR^ ej2^CR^/+* heterozygotes exhibit moderate branching similar to the natural mutant, revealing a clear gene dosage effect. This dosage-sensitive genetic interaction can be exploited for quantitative manipulation of inflorescence branching by altering *EJ2* dosage in *j2* null mutant backgrounds. Most notably, *j2^TE^ ej2^W^/+* hybrids display mildly branched inflorescences with more flowers, conferring a heterotic effect for fruit yield (23).

Previously, it has been proposed that dosage-sensitive interactions can be exploited to induce quantitative variation by CRISPR-Cas genome editing (25, 26). However, the stability of gene dosage relationships across genetic backgrounds remains largely unexplored. Here, we show that cryptic variation modulates the phenotypic outcome of gene dosage changes, presenting a previously unrecognized layer of genetic complexity that can limit the predictability of genome editing for crop improvement.

## Results

Previous work demonstrated that reducing the number of functional gene copies of *J2* and *EJ2* leads to gradual increases in inflorescence branching, showing that tomato inflorescence architecture is sensitive to changes in MADS-box gene dosage (6, 23). To examine whether this dosage relationship is background-dependent, we generated *j2^null^* and *ej2^null^* alleles in two genetically distinct tomato accessions, the domestic cultivar Sweet-100 (*S. lycopersicum* cv. S100) and the wild ancestor species *S. pimpinellifolium* (acc. LA1589), using CRISPR-Cas genome editing (**Fig. 1a,b**). As expected, *j2^null^ ej2^null^*double mutants developed strongly branched inflorescences with more than 30 branching events per inflorescence in both domesticated and wild tomato whereas the single mutants produced unbranched inflorescences (**Fig. 1 c–f**). Thus, the genetic interaction between *J2* and *EJ2* genes is conserved between wild and domesticated tomato. To investigate the effect of genetic background on MADS-box dosage sensitivity, we analyzed plants with different combinations of homozygous and heterozygous *j2^null^* and *ej2^null^* mutations in both wild and domesticated genotypes. Interestingly, while plants homozygous for *j2^null^* and heterozygous for *ej2^null^* (*j2^null^ ej2^null^/+*) displayed strong branching (∼11.9 branching events) in the domesticated background, we observed only very mild branching (∼0.4 branching events) in the wild accession (**Fig. 1 c–f**). Similarly, *j2^null^/+ ej2^null^* mutants developed weakly branched inflorescences in the domesticated background while branching was fully suppressed in wild tomato (**Fig. 1 c–f**). These observations provided first evidence that MADS-box gene dosage sensitivity differs between distinct wild and domesticated backgrounds.

**Fig. 1.**
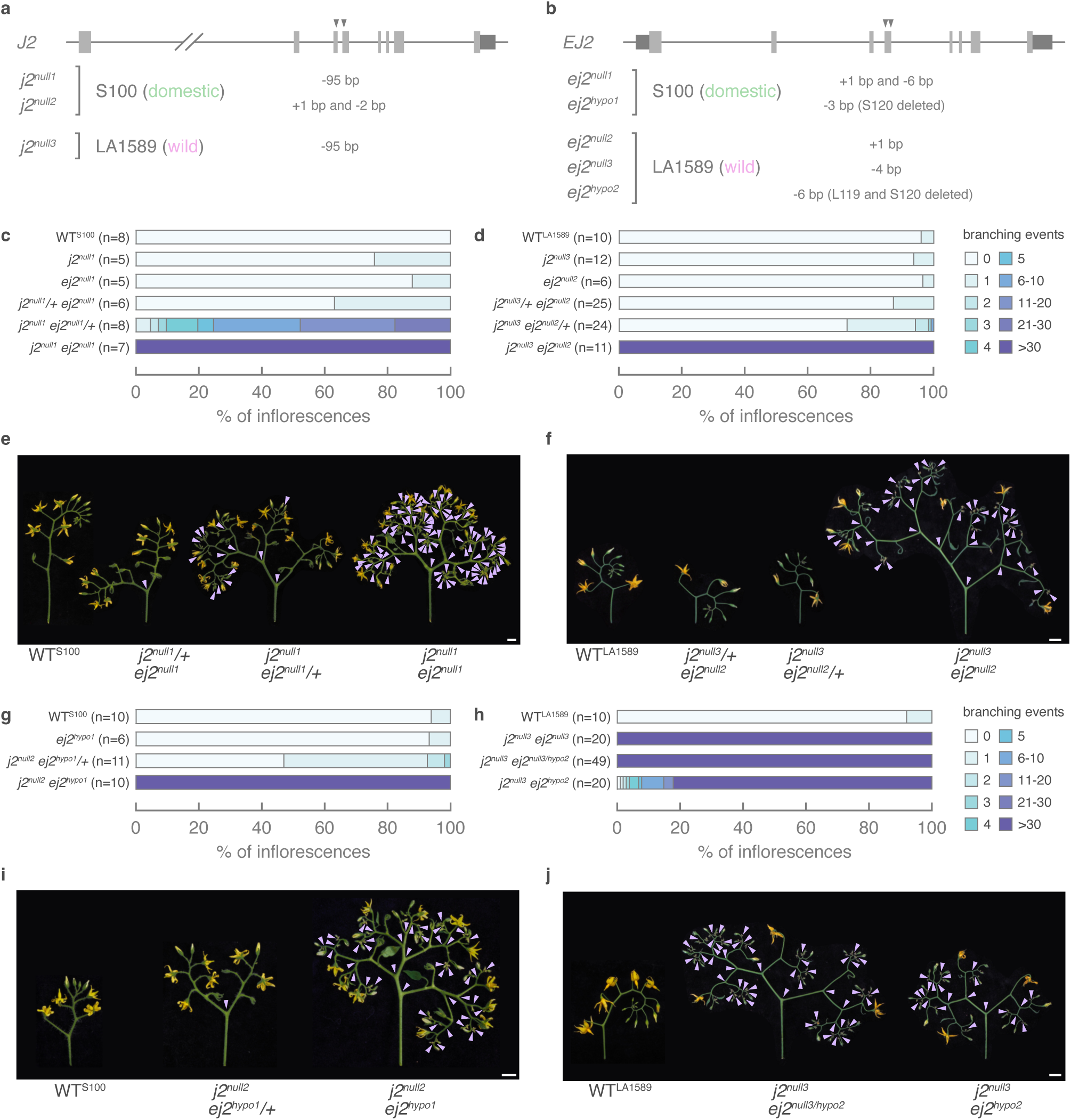
MADS-box dosage sensitivity differs between domesticated and wild tomato. **a**,**b**, Schematic representation of *J2* (**a**) and *EJ2* (**b**) with location of the CRISPR-Cas9 target sites and mutant alleles in domestic (*S. lycopersicum* acc. S100) and wild (*S. pimpinellifolium* acc. LA1589) tomato. **c**,**d**, Quantification of inflorescence branching in *j2^null^ ej2^null^* segregating populations in domestic (**c**) and wild (**d**) tomato. **e**,**f**, Representative images of domestic (**e**) and wild (**f**) inflorescences with different strengths of branching from genotypes segregating *j2^null^ ej2^null^* alleles. **g**,**h**, Quantification of inflorescence branching in *j2^null^ ej2^null/hypo^* segregating populations in domestic (**g**) and wild (**h**) tomato. **i**,**j**, Representative images of domestic (**i**) and wild (**j**) inflorescences with different strengths of branching from genotypes segregating *j2^null^ ej2^null/hypo^* alleles. Gene models in **a** and **b**: exons, untranslated regions, and Cas9 cleavage sites for guide RNAs are indicated by light gray boxes, dark gray boxes, and gray arrowheads, respectively. Per genotype, the number of individual plants for which 5 inflorescences were counted in **c**, **d**, **g**, and **h** is indicated by *n*. Scalebars and arrowheads in **e**, **f**, **i**, and **j** represent 1 cm and indicate inflorescence branching events, respectively. *J2*, *JOINTLESS2*; *EJ2*, *ENHANCER OF J2*; WT, wild-type.

To further support this finding, we exploited partial loss-of-function (hypomorphic) *ej2^hypo^* alleles that lead to the loss of amino acid residues in the K-domain, which is responsible for MADS-box oligomerization (27, 28). In Arabidopsis (*A. thaliana*), mutations in the K-domain of SEP3 have been shown to partially reduce *SEP3* activity (29, 30). We obtained *ej2^hypo^* alleles that lead to the loss of the K-domain residues serine-120 (S120) and leucine-119/serine-120 (L119/S120) in domestic and wild tomato, respectively (**Fig. 1b**). In Arabidopsis SEP3, the corresponding leucine and serine residues are in a kink region that orients the two α-helices of the K-domain at an ∼90° angle (31), and mutating the leucine residue decreased tetramer formation (32). We modelled the structure of tomato EJ2 homotetramers using AlphaFold3, which indicated that deletion of L119 and S120 substantially reduces the kink angle and likely impacts tetramer functionality (**Fig. S1a–c**). Indeed, *ej2^hypo^*homozygosity consistently conferred strong branching in a domestic *j2^null^*background, demonstrating that deletion of L119/S120 impacts EJ2 function (**Fig. 1g–j**). However, inflorescence branching was more variable in wild tomato, with 18.0% of inflorescences showing suppression of branching at different degrees (**Fig. 1h**). Taken together, these results show that inflorescence architecture of wild tomato is less sensitive to reductions in MADS-box gene dosage compared with domestic tomato. This suggests that the dosage-sensitive genetic interaction between these MADS-box genes is modified by cryptic variant(s) in wild tomato.

To identify the genetic basis of variation in MADS-box dosage sensitivity between domestic and wild tomato, we utilized a previously published recombinant F_2_ population derived from a cross between a natural hypomorphic *j2^TE^ ej2^W^* mutant (*compound inflorescence2* or *s2)* in domestic tomato and the wild tomato accession LA1589 (**Fig. S2a,b**) (23). The natural *s2* mutant develops branched inflorescences due to a transposon insertion in *J2* that eliminates its function and an intronic insertion in *EJ2* that partially reduces functional *EJ2* transcript levels (23). Approximately 24% of the *j2^TE^ ej2^W^* double mutant plants from the F_2_ population displayed suppression of inflorescence branching at various levels, consistent with the presence of cryptic variant(s) in wild tomato that modify inflorescence architecture (**Fig. S2c**) (23). To investigate this cryptic suppression of inflorescence branching, we further introgressed the *j2^TE^ ej2^W^* mutations from *s2* into wild tomato. After two rounds of backcrossing, we obtained *j2^TE^ ej2^W^* individuals that displayed complete suppression of inflorescence branching (**Fig. S2a,d,e**), indicating that wild tomato indeed harbors cryptic modifiers of inflorescence architecture.

Next, we re-analyzed published QTL sequencing data for the recombinant population and mapped two major QTLs on chromosome 1 and 2 (**Fig. S3 and Fig. 2a,b**). We identified the previously characterized *suppressor of branching 1* (*sb1*) locus on chromosome 1, which contains a copy number variant of the MADS-box gene *SISTER OF TM3* (*STM3*). The single-copy *STM3* haplotype partially suppresses inflorescence branching and mitigates the negative effect of *j2^TE^ ej2^W^*mutations on fruit productivity in modern tomato genotypes (**Fig. 2a,b**) (6, 33). In addition, we identified a second, uncharacterized large-effect locus that spans a region of ∼7.6 Mb on chromosome 2, which we designated *suppressor of branching 2* (*sb2*) (**Fig. 2a,b**) (23). We fine-mapped *sb2* to an interval of ∼160 Kb using linkage analysis and recombinant inbred lines that we isolated from subsequent generations of *sb2* segregating populations (**Fig. 2c and Table S1–4**). To separate effects from *sb1*, we conducted fine-mapping in segregating populations that were homozygous for the suppressing *sb1* haplotype. Using recombinant genotypes 3 and 4 of our fifth fine-mapping population, we finally resolved *sb2* to a region of ∼83 Kb containing 15 annotated genes (**Fig. 2d–f**).

**Fig. 2.**
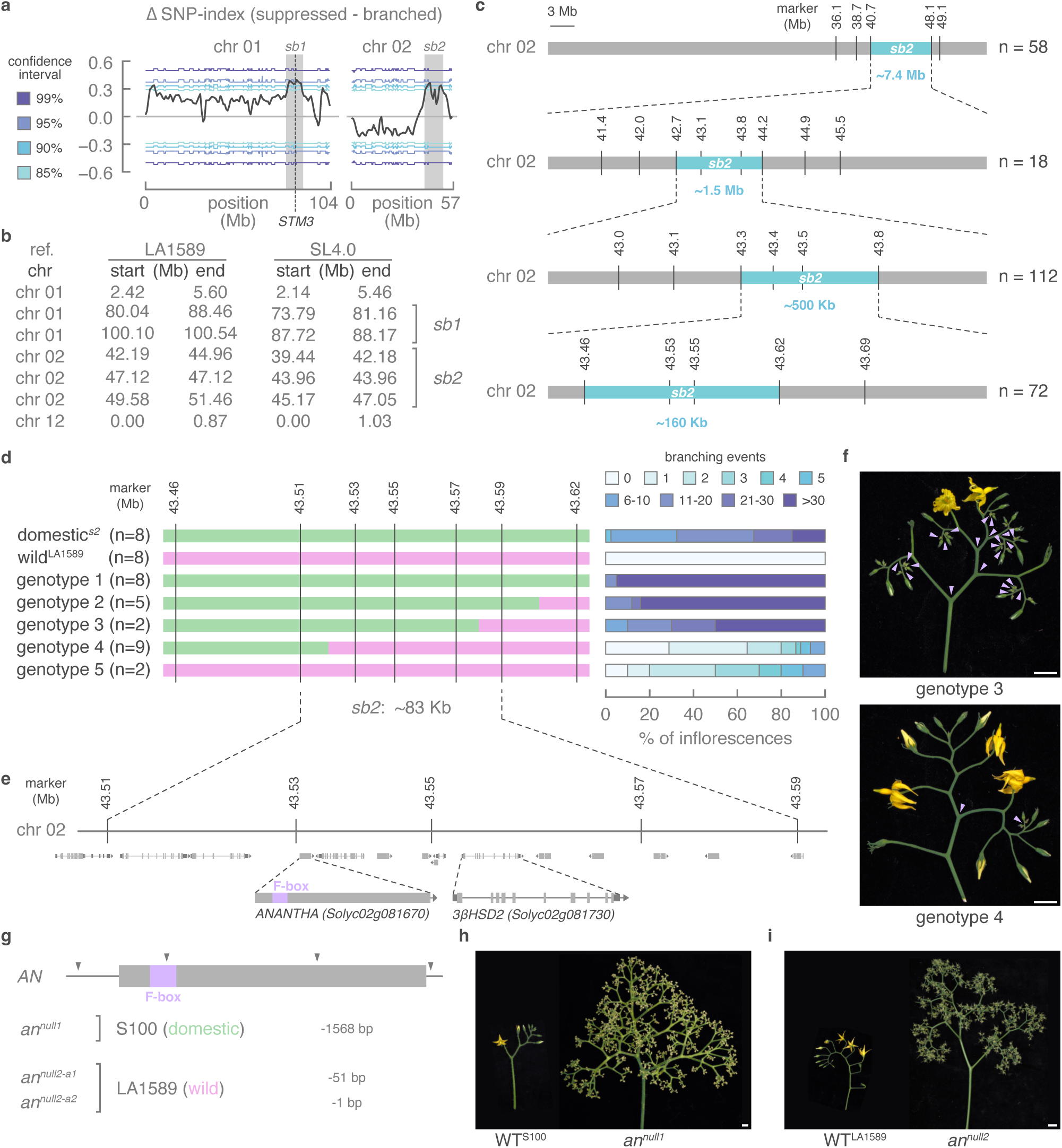
Suppression of branching in wild tomato maps to a region of ∼83 Kb containing *ANANTHA* and *3βHSD2*. **a**, Quantitative trait locus (QTL) sequencing using bulked segregants of plants with branched and suppressed inflorescences from a *s2* x LA1589 F2 population showed two *suppressor of branching* (*sb*) loci from wild tomato (acc. LA1589) on chromosomes 1 and 2. The *sb1* QTL contains a copy number variant of the MADS-box gene *SISTER OF TM3* (*STM3*). **b**, Physical positions of the QTL regions shown in (**a**) in wild (LA1589) and domestic (SL4.0) tomato. **c**, The *sb2* locus was fine-mapped to an interval of ∼160 Kb between markers 43.46 Mb and 43.62 Mb (SL4.0 coordinates) on chromosome 2 in subsequent generations of *sb2* segregating populations. Blue boxes indicate the fine-mapped *sb2* locus in each population. **d**, The *sb2* locus was fine-mapped to an interval of ∼83 Kb between markers 43.51 Mb and 43.59 Mb (SL4.0 coordinates) on chromosome 2. Each distinctive genotype is represented by a horizontal bar. Green and pink shading indicates homozygosity for *s2* and LA1589, respectively. Per genotype, the number of individual plants for which 5 inflorescences were counted is indicated by *n*. **e**, Gene models and coordinates of the 15 genes in the *sb2* locus including *ANANTHA* and *3βHSD2*. **f**, Representative images of strongly branched and suppressed inflorescences from plants with genotype 3 and 4 from (**d**), respectively. **g**, Schematic representation of *ANANTHA* with location of the CRISPR-Cas9 target sites and mutant alleles in domestic (*S. lycopersicum* acc. S100) tomato. Gene model: exon, region encoding F-box, and Cas9 cleavage sites for guide RNAs are indicated by a light gray box, a lavender box, and gray arrowheads, respectively. **g**,**h**, Representative images of cauliflower inflorescences from *anantha* mutants in domestic (acc. S100) (**h**) and wild (acc. LA1589) (**i**) tomato. Scalebars and arrowheads in **f**, **h**, and **i** represent 1 cm and indicate inflorescence branching events, respectively. WT, wild-type; *AN*, *ANANTHA*; *3βHSD2*, *3β-hydroxysteroid dehydrogenase/C4-decarboxylase 2*.

To identify the causative gene(s) underlying *sb2*, we first surveyed all genes in this region for coding sequence variants. Of the 15 genes, four carried missense mutations and one contained exonic in-frame insertions, yet none of these five genes are expressed in meristem tissues (**Table S5 and Fig. S4**) (19). Among the eight genes that are expressed in meristems, only *ANANTHA* (*AN*) is dynamically expressed during meristem maturation with an expression peak at the floral stage (**Fig. S4**). The *AN* gene encodes an F-box protein that is orthologous to Arabidopsis UNUSUAL FLORAL ORGANS (UFO), which forms a floral specification complex together with transcription factor LEAFY (34, 35). Loss of *AN* activity prevents meristems from adopting floral fate and results in excessive inflorescence branching in domestic tomato (**Fig. 2g,h**) (36–38), and we observed a comparable phenotype in wild tomato *an* null mutants that were generated for the purpose of this study (**Fig. S5a and Fig. 2g,i**).

Absence of any *AN* coding variants between the domestic *s2* mutant and the wild accession LA1589 suggested regulatory variant(s) to be underlying suppression of inflorescence branching by the *sb2* locus. To identify genetic variants that may alter *AN* transcriptional regulation, we scanned noncoding regions up- and downstream of the *AN* gene for sequence variation between the domestic *s2* (or *j2^TE^ ej2^W^*) mutant and wild accession LA1589 (**Fig. S5b**). The only region of accessible chromatin in meristem tissue upstream of *AN*, which overlaps with a cluster of conserved non-coding sequences (CNSs) and whose removal by CRISPR-Cas genome editing led to inflorescence branching (39, 40), did not contain any genetic variants (**Fig. S5b**). We did detect a downstream SNP within a CNS, and a 1-bp insertion/deletion (InDel) overlapping with a predicted DNA binding with one zinc finger (DOF) binding site but positioned adjacent to the core AAAG motif (**Fig. S5b**). In addition, we identified an 8-bp InDel located 457 bp upstream of the *AN* start codon, which partially overlaps with a predicted APETALA2/Ethylene Responsive Factor (AP2/ERF) binding site (**Fig. S5b**) (41, 42). Several AP2/ERFs have already been implicated in the control of organ development at the shoot apical meristem in tomato and other species (43–46). AP2/ERFs often act as transcriptional repressors through an ETHYLENE RESPONSE FACTOR (ERF)-ASSOCIATED AMPHIPHILIC REPRESSION (EAR) motif (47) that allows them to recruit co-repressors of the TOPLESS family (48, 49). We reasoned that disruption of the predicted AP2/ERF binding site in the wild tomato accession could explain suppression of inflorescence branching by the *sb2* locus. To test this hypothesis, we selected a recombinant inbred line with branched inflorescences that was homozygous for the domestic *SB2* haplotype and used CRISPR genome editing to disrupt the 8-bp insertion containing the AP2/ERF binding motif **(Fig. S5c,d)**. We isolated six independent *an^reg^* alleles in the T1 generation, of which five had InDels disrupting the motif and one that removed the motif completely (**Fig. S5c**). However, all T1 plants with *an^reg^* alleles produced inflorescences that displayed strong branching similar to the parental *SB2* line (**Fig. S5e,f**), suggesting that the 8-bp InDel is not the causative variant in the *sb2* locus.

In addition, we conducted bulk RNA sequencing on dissected meristems at the transition and floral stage of meristem maturation (19) from plants harboring the domestic *SB2* and wild *sb2* haplotype. We found 329 and 278 differentially expressed genes (DEGs, |log_2_FC| ≥ 0.585, FDR<0.05) between genotypes at transition and floral meristem stages, respectively (**Fig. 3a,b, Table S6,7**). We performed hierarchical clustering of DEGs followed by gene ontology (GO) enrichment analyses of clusters with similar expression patterns and found that cluster 1, which was downregulated in the suppressed *sb2* genotype, was enriched for terms related to sterol metabolic processes (**Fig. 3b–d**). Among these were key cholesterogenesis genes including *STEROL SIDE CHAIN REDUCTASE 2* (*SSR2*) and *STEROL C-5*(*6*) *DESATURASE 2* (*C5-SD2*) (50, 51), as well as orthologs of Arabidopsis genes required for brassinosteroid biosynthesis including *ROTUNDIFOLIA 3* (*ROT3*) (**Fig. 3e,f**) (52). Surprisingly, we did not detect any significant changes in *AN* transcript levels between genotypes at the transition or floral meristem stage (**Fig. 3g,h, Fig. S6a**). Among all meristem-expressed genes at the *sb2* locus, we only observed a modest (1.39-fold), but significant (FDR=0.018) decrease in the expression of sterol biosynthesis gene *3β-hydroxysteroid dehydrogenase/C4-decarboxylase 2* (*3βHSD2*) (51) in the suppressed *sb2* genotype at the transition stage (**Fig. 3g**). The *3βHSD2* gene encodes an enzyme involved in the biosynthesis of cholesterol and phytosterols, the respective precursors of steroidal glycoalkaloids and brassinosteroid hormones (51). Using quantitative PCR, we verified the absence of a significant *AN* expression change, but we could not validate the decrease in *3βHSD2* expression in *sb2* transition meristems (**Fig. S6b)**. Despite this discrepancy with our RNA-seq results, we hypothesized that lower *3βHSD2* expression from *sb2* may cause the downregulation of other phytosterol biosynthesis genes and thereby suppresses inflorescence branching. To functionally test if reduced *3βHSD2* activity affects inflorescence branching, we generated two independent *3bhsd2* knockout alleles by CRISPR-Cas genome editing (**Fig. 3i and Fig. S7a**). However, loss of *3βHSD2* activity did not alter inflorescence architecture in *3bhsd2* single mutants nor suppressed inflorescence branching in isogenic *j2^null2^ej2^hypo1^ 3bhsd2* triple mutants (**Fig. 3j,k and Fig. S7b**). We also tested if reduced *3βHSD2* activity suppresses MADS-box dosage effects on inflorescence architecture that we had observed in the *j2^null2^ej2^hypo1^/+* genotype (see **Fig. 1g,i**). Although *j2^null2^ej2^hypo1^/+* consistently displayed weak inflorescence branching, the dosage effect was not suppressed by either homozygous or heterozygous *3bhsd2* mutations in *j2^null2^ej2^hypo1^/+ 3bhsd2* and *j2^null2^ej2^hypo1^/+ 3bhsd2/+* genotypes (**Fig. 3j,k** and **Fig. S7b**). We concluded that the *3βHSD2* expression change in *sb2* is not causative for the suppression of inflorescence branching. Given the conserved function of *AN* as a floral identity gene, we speculate that variation at *sb2* affects the spatiotemporal pattern of *AN* expression rather than its absolute expression level, and thus causes the suppression of inflorescence branching in wild tomato.

**Fig. 3.**
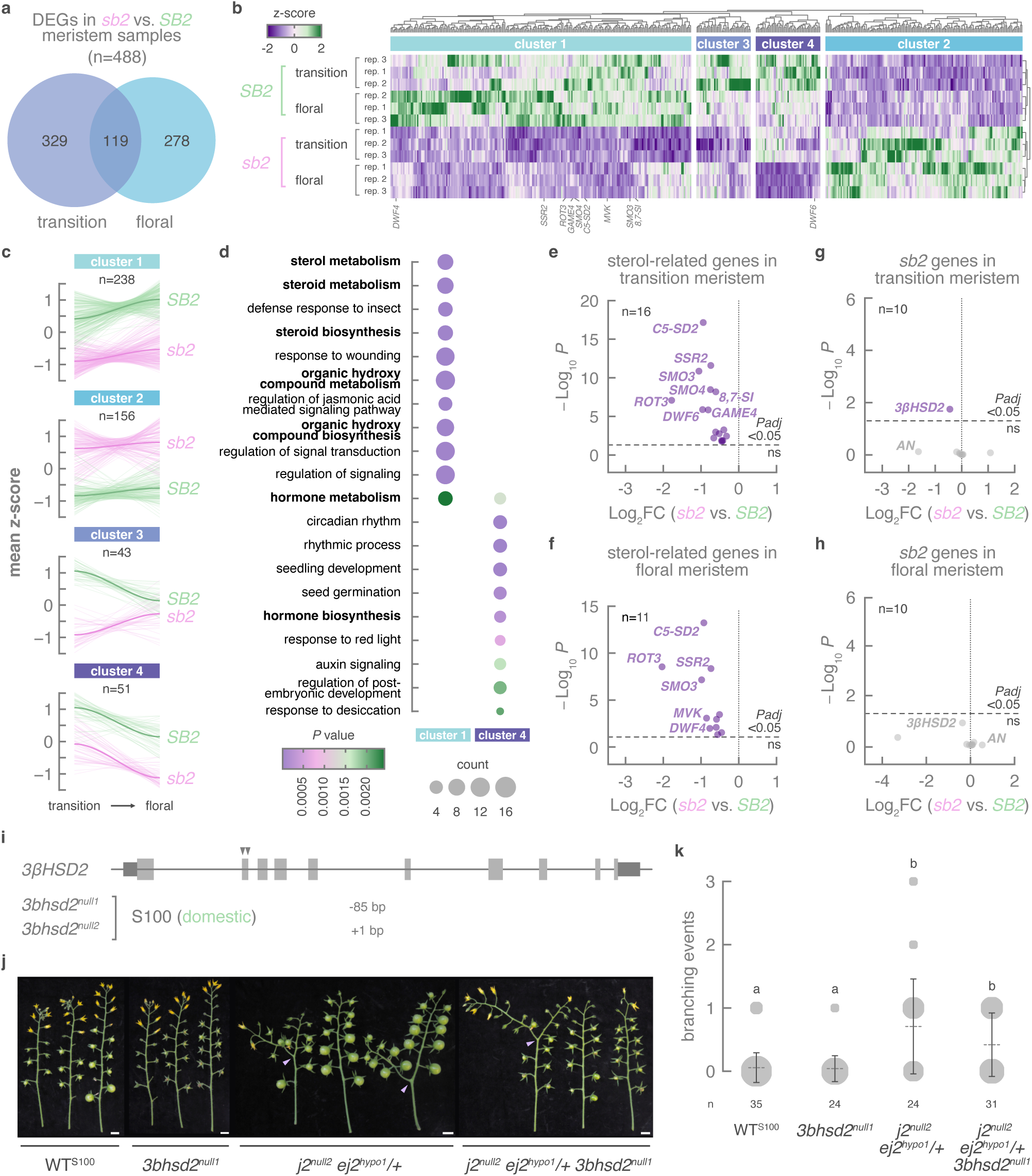
Suppression of branching by *sb2* leads to changes in steroid-related gene expression. **a**, Number of differentially expressed genes (DEGs; log₂ fold change |≥| 0.585 and FDR ≤ 0.05) between plants harboring the domestic *SB2* and wild *sb2* haplotype, based on RNA sequencing of dissected meristems at the transition and floral stage. **b**, Heatmap depicting z-score normalized expression of DEGs (n=488) between *SB2* and *sb2* plants. **c**, Z-score normalized expression profiles of DEG clusters with similar expression patterns identified by hierarchical clustering in (**b**). **d**, The 10 most enriched gene ontology (GO) categories for DEGs in clusters 1 and 4. No GO enrichment was detected for clusters 2 and 3. *P* values were obtained using the Benjamini-Hochberg (BH) method in clusterProfiler. **e**,**f** Volcano plots displaying downregulation of sterol-related genes in transition (**e**) and floral (**f**) meristems of *sb2* compared with *SB2* plants. **g**,**h**, Volcano plots displaying expression patterns for genes at the *sb2* locus between *SB2* and *sb2* plants in transition (**g**) and floral (**h**) meristems. **i**, Schematic representation of *3βHSD2* with location of the CRISPR-Cas9 target sites and mutant alleles in domestic (*S. lycopersicum* acc. S100) tomato. Gene model: exons, untranslated regions, and Cas9 cleavage sites for guide RNAs are indicated by light gray boxes, dark gray boxes, and gray arrowheads, respectively. **j**, Representative images of inflorescences from *3bhsd2^null1^*, *j2^null2^ ej2^hypo1^/+,* and *j2^null2^ ej2^hypo1^/+ 3bhsd2^null1^* plants. Scalebars and arrowheads represent 1 cm and indicate inflorescence branching events, respectively. **k**, Quantification of inflorescence branching for *3bhsd2^null1^*, *j2^null2^ ej2^hypo1^/+,* and *j2^null2^ ej2^hypo1^/+ 3bhsd2^null1^* plants. Dotted lines represent mean number of branching events for each genotype. Error bars denote standard deviation (*n*=24–35). Circle areas represent the number of inflorescences per genotype. Statistical significance was determined by ANOVA followed by Tukey’s post-hoc analysis (*P* < 0.05; indicated by different letters). *suppressor of branching 2, sb2; DWF*, *DWARF*; *SSR2*, *STEROL SIDE CHAIN REDUCTASE 2*; *ROT3*, *ROTUNDIFOLIA 3*; *GAME4*, *GLYCOALKALOID METABOLISM 4*; *SMO*, *C-4 STEROL METHYL OXIDASE*; *C5-SD2*, *STEROL C-5*(*6*) *DESATURASE 2*; *MVK*, *MEVALONATE KINASE*; *8,7-SI*, *STEROL 8,7 ISOMERASE*; *3βHSD2*, *3β-hydroxysteroid dehydrogenase/C4-decarboxylase 2*; *AN*, *ANANTHA; J2*, *JOINTLESS2*; *EJ2*, *ENHANCER OF J2*; WT, wild-type.

**Fig. 4.**
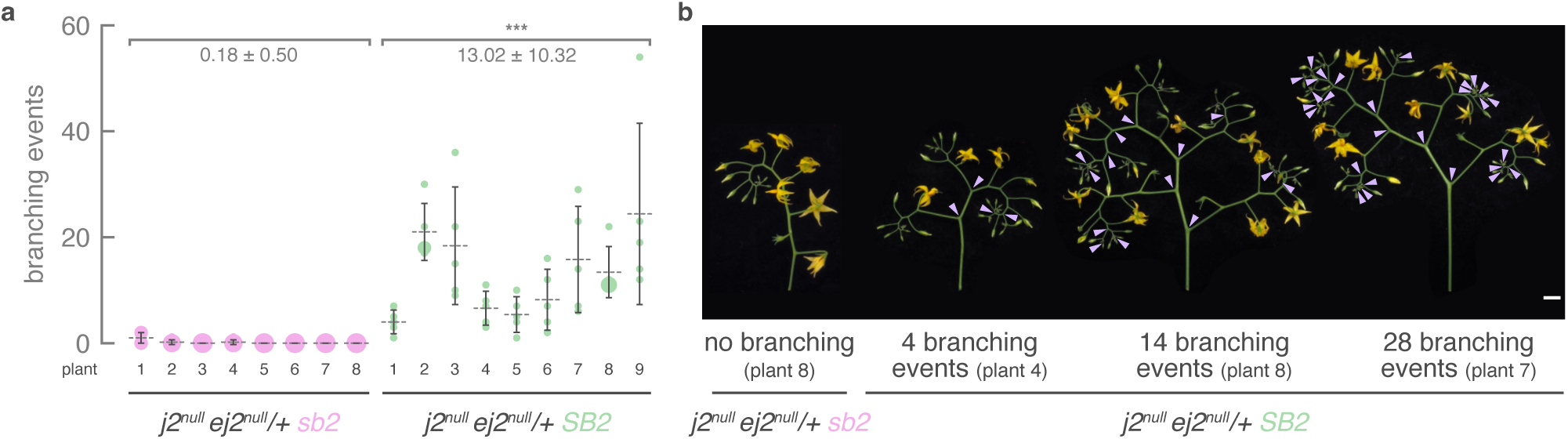
The natural *sb2* haplotype suppresses MADS-box dosage sensitivity in CRISPR-engineered *j2 ej2/+* genotypes. **a**, Quantification of inflorescence branching for *j2^null^ ej2^null^/+* hybrids that are homozygous for the wild *sb2* or domestic *SB2* haplotype. Dotted lines represent mean number of branching events for each individual plant. Error bars denote standard deviation (*n*=5). Circle areas represent the number of inflorescences for each *j2^null^ ej2^null^/+ sb2* (pink circles) and *j2^null^ ej2^null^/+ SB2* (green circles) plant. **b**, Representative images of inflorescences with different strengths of branching from *j2^null^ ej2^null^/+ sb2* and *j2^null^ ej2^null^/+ SB2* plants. The number of branching events on each imaged inflorescence is specified. Scalebars and arrowheads represent 1 cm and indicate inflorescence branching events, respectively. *suppressor of branching 2, sb2*; *J2*, *JOINTLESS2*; *EJ2*, *ENHANCER OF J2*.

Finally, we tested whether cryptic variation at *sb2* can be leveraged to improve the predictability of CRISPR-engineered MADS-box dosage effects on inflorescence branching. The distinct branching phenotypes of genome-edited MADS-box mutants in domestic and wild backgrounds demonstrated that inflorescence architecture of wild tomato is less sensitive to engineered dosage effects compared with domestic tomato (**Fig. 1** and **Fig. S2d,e**). To test whether those differences in MADS-box dosage sensitivity are due to *sb2*, we generated a recombinant population that segregated both *sb2* and the genome-edited *j2^null^* and *ej2^null^* alleles in a homozygous *sb1* background (**Fig. S2f**). Since MADS-box dosage effects on inflorescence branching are most strongly expressed in mutants homozygous for *j2^null^* and heterozygous for *ej2^null^* (**Fig. 1c–f**), we isolated *j2^null^ ej2^null^/+* mutants that were either homozygous for the wild *sb2* or domestic *SB2* allele (**Fig. S2f**). While plants harboring the wild *sb2* allele showed almost complete suppression of branching, those with the domestic *SB2* allele regained the capacity to branch, producing inflorescences with an average of 4.0 to 24.4 branching events per plant (**Fig. 3a,b**). The broad phenotypic variation between *SB2* individuals indicates that additional minor-effect QTLs are involved in the branching suppression. Taken together, our results demonstrate that cryptic variation at *sb2* exposes dosage sensitivity of both naturally occurring and genome-edited MADS-box mutations.

## Discussion

Leveraging dosage-sensitive systems to fine-tune quantitative traits using genome editing holds enormous potential for crop breeding. However, genetic interactions between dosage-sensitive alleles and cryptic variants in distinct genetic backgrounds can result in unexpected, and sometimes undesirable, phenotypic outcomes. Here, we discovered that dosage-sensitive MADS-box mutations that can be exploited to optimize inflorescence architecture for fruit yield (23) behave differently in distinct wild and domesticated tomato genomes. We showed that reductions in dosage of the MADS-box genes *J2* and *EJ2*, either through null or hypomorphic alleles, leads to a weaker effect on inflorescence branching in wild compared with domestic tomato. We fine-mapped this difference to cryptic variation at the *sb2* locus, which comprises a region of ∼83 Kb on chromosome 2 and contains *AN*, a well-known regulator of tomato inflorescence architecture (36, 37). We did not identify any CDS variants or gene expression changes for *AN* in meristems of plants harboring the wild *sb2* and domestic *SB2* allele. We speculate that regulatory sequence variation at *sb2* leads to changes in spatiotemporal *AN* expression pattern, explaining the suppression of inflorescence branching. A similar scenario underlies species-specific differences in flower organ (petal) number variation between Arabidopsis and its close relative *Cardamine hirsuta*, where expression of the MADS-box gene *APETALA1* is restricted to fewer cells in *C. hirsuta* floral meristems, yielding a variable petal number in contrast with a robust pattern of four petals in Arabidopsis (53). Although we did not fully resolve *sb2* genetically, we show *sb2* can be utilized to suppress inflorescence branching from both natural and engineered *j2* and *ej2* CRISPR alleles in wild tomato. In addition to *sb2*, wild tomato harbors *sb1*, a previously characterized copy number variant of the MADS-box gene *STM3* that partially suppresses inflorescence branching in domestic *j2^TE^ ej2^W^* mutants (6, 33). The domestic cultivar (Sweet-100) in which we engineered *J2*/*EJ2* dosage effects carries neither *sb1* nor *sb2*. Therefore, differences in MADS-box dosage sensitivity between wild and domestic tomato likely reflect the combined effect of *sb1* and *sb2*, which may act additively or synergistically, and we obtained evidence for additional minor-effect QTLs to modulate inflorescence architecture. Together, these findings indicate that inflorescence architecture in tomato is controlled by multiple cryptic modifiers that complicate the phenotypic outcomes from MADS-box dosage changes.

The transcription factors encoded by the dosage-sensitive MADS-box genes, *J2* and *EJ2*, are known to act in tetrameric complexes to regulate the activity of shoot apical meristems (22, 23). Although they exhibit redundant roles during inflorescence development, stronger branching was observed for *j2^null^ ej2^null^/+* compared with *j2^null^/+ ej2^null^*heterozygotes in both domestic and wild tomato. This asymmetry in dosage effect may reflect differences in (post)-transcriptional and translational buffering of *J2* and *EJ2* (*54*), reinforcing the gene balance hypothesis, which posits that changes in relative abundance of complex members can affect complex activity and downstream developmental outcomes (10). The relevance of the gene balance hypothesis for MADS-box function is further supported by prior work showing that a tandem duplication of the hypomorphic *ej2^W^* allele suppresses inflorescence branching in domestic *j2 ej2^W^* mutants (6). Our findings extend the gene balance hypothesis by demonstrating that phenotypic consequences from gene dosage perturbation depend on the genotypic context. Thus, the architecture of gene dosage sensitivity can evolve and our discovery of *sb2* as a cryptic modifier reveals how hidden genetic variation shapes this process. While cryptic variants can contribute to the evolvability of developmental networks in natural populations (8), they can also constrain predictable outcomes from precision breeding of quantitative traits by genome editing. The phenotypic outcome of targeted, engineered mutations in dosage-sensitive systems can therefore not be fully predicted without considering the broader genetic context in which they occur. Knowledge of the allelic state of known and still unknown modifiers of inflorescence architecture could inform the design of edits to achieve the desired level of inflorescence complexity to optimize fruit yield in specific genotypes. As precise engineering of protein coding sequences and gene regulatory regions at multiple loci becomes a new routine, information on cryptic modifiers will be critical to transform genome editing from a descriptive research approach to a predictive breeding approach for engineering complex traits in crops.

## Materials and Methods

### Plant material and growth conditions

Seeds of *S. lycopersicum* cv. Sweet-100 (S100) double-determinate (38), S100 *j2^null1^ ej2^null1^* mutant (55), S100 *j2^null2^ ej2^hypo1^* mutant (56), S100 *anantha* mutant (38), *S. pimpinellifolium* acc. LA1589, and *compound inflorescence 2* (*s2*; LA4371) were sown from our own stocks. Seeds were either pre-germinated on moistened Whatman paper at 28°C in complete darkness or directly sown and germinated in soil in 96-cell plastic flats. For greenhouse experiments, plants were grown under long-day photoperiods (16:8 h) under natural light supplemented with artificial light from high-pressure sodium bulbs or LED panels (∼200 μmol m^−2^ s^−1^), under constant temperature of 25°C and relative humidity of 50**–**60%. Tomato plants were grown in 5 l pots (two plants per pot) under drip irrigation and standard fertilizer regimes. For field experiments, plants were grown in open fields at Cold Spring Harbor Laboratory, Cold Spring Harbor, New York, US or in open fields or polytunnel at the University of Lausanne, Lausanne, Vaud, Switzerland.

### CRISPR-Cas cloning

CRISPR-Cas9 mutagenesis in was performed as previously described (38, 55). Briefly, guide RNAs (gRNAs) were designed using CRISPOR and the Sweet-100v.2.0 (38) and the LA1589v0.1 (57)reference genomes. Binary vectors for CRISPR-Cas mutagenesis were assembled using the Golden Gate cloning system as previously described (38, 58, 59). For targeting *J2* and *EJ2*, EJ2-sgRNA-1, EJ2-sgRNA-2, J2-sgRNA-1, and J2-sgRNA-2 were cloned into Level(L)1 acceptors pICH47751, pICH47761, pICH47772, pICH47781, respectively, and combined into the L2 acceptor pICSL4723-P1 (addgene #86173) with pICSL11024 (addgene #51144, Nos::NptII::ocs), pICH47742_2×35S::SpCas9-P2A-GFP::nosT (57), and pICH41822 to obtain pSS009. We transformed pSS009 into S100 to generate *j2^null2^*and *ej2^hypo1^* alleles and into LA1589 to obtain *j2^null3^*, *ej2^null2^*, *ej2^null3^*, *ej2^hypo2^* alleles. For targeting the coding sequence of *AN*, AN-CDS-sgRNA-1**–**4 were cloned into L1 acceptors pICH47751, pICH47761, pICH47772, pICH47781, respectively, and combined into pICSL4723-P1 with pICSL11024, pSS082 (55), and pICH41822 to obtain pSS123. pSS123 was transformed into LA1589. For targeting upstream noncoding sequences of *AN*, a new L1 part pICH47742_pSlUBQ10:SpRY-P2A-GFP (pSS121) was cloned. The coding sequence of SpRY was Golden Gate domesticated by amplifying four fragments, i.e. PCR1 (P584+P577), PCR2 (P578+P579), PCR3 (P580+P581), and PCR4 (P582+P583) from the template pYPQ166-SpRY (addgene #161520) (60)PCR1 to PCR4 were individually cloned into the universal L-1 acceptor pAGM1311 and combined in the L0 acceptor pAGM1287 to generate pSS060, which was combined with pAGM1301_P2A-GFP (57), pSS051 (55) and pICH41421 in the L1 acceptor pICH47742 to obtain pSS120 (pSlUBQ10:SpRY-P2A-GFP). AN-CRE-sgRNA-1**–**3 were cloned into L1 acceptors pICH47761, pICH47772, pICH47781, respectively, and combined into the L2 acceptor pICSL4723-P1 with pICSL11024, pSS120, pLL034 (PcUBQ-SlGRF-GIF) (55), and pICH41822 to obtain pSS121. To mutate *3βHSD2*, 3bHSD2-sgRNA-1 and 3bHSD2-sgRNA-2 were cloned into pICH47751 and pICH47761, respectively, and combined in pICSL4723-P1 with pICSL11024, pICH47742_2×35S::SpCas9-P2A-GFP::nosT (57), and pICH41780 to obtain pSST039, which was transformed pSST039 into S100. All primers and gRNAs used in this study are listed in **Table S8** and **S9**, respectively. All CRISPR-Cas mutant alleles used in this study, including those generated with these constructs and those obtained from previous work, are listed in **Table S10**.

### Plant transformation

Binary vectors were transformed into domesticated tomato (*S. lycopersicum* cv. S100), wild tomato (*S. pimpinellifolium* acc. LA1589), and a recombinant inbred line from the first *sb2* fine-mapping population that was homozygous for the domestic *s2* parent at the *SB2* locus by *Agrobacterium*-mediated transformation according to Gupta and Van Eck (61)with minor modifications as described in (55).

### Plant genotyping

Genomic DNA was prepared from homogenized leaf tissue using an extraction buffer (pH 9.5) containing 0.1 M of tris(hydroxymethyl)aminomethane-hydrochloride (Tris-HCl), 0.25 M of potassium chloride (KCl), and 0.01 M of ethylenediaminetetraacetic acid (EDTA). This mixture was incubated at 95°C for 10 min and subsequently cooled at 4°C for 5 min. After addition of 3% (w/v) BSA, collected supernatant was used as a template for PCR. All primers used for genotyping are listed in **Table S8**.

#### Identification of CRISPR-Cas mutants

To identify CRISPR-Cas *j2^null^ ej2^null/hypo^*, *anantha*, and *an^reg^* T0 mutants, gRNA target regions were amplified with gene-specific primers (**Table S8**). The PCR amplicons were either subjected to amplicon deep sequencing, or they were purified using ExoSAP-IT (Thermo Fisher Scientific) and analyzed by Sanger sequencing of the purified PCR amplicons, followed by decomposition of quantitative sequence trace data using Inference of CRISPR Editing (ICE) CRISPR Analysis Tool (https://ice.synthego.com/#/). In subsequent generations, mutant alleles were identified using the same approaches, based on gel lesions, by using single-tube markers (62) or by using cleaved amplified polymorphic sequence (CAPS) markers (**Table S8**).

Amplicon deep sequencing was performed for high throughput genotyping as previously described (63) with minor modifications. Briefly, gRNA target regions were amplified using gene-specific primers (**Table S8**) with common adapter sequences. Gene-specific amplicons were diluted ten-fold in H_2_O before they were used as template for the addition of sample-specific indexes using adapter primers with unique barcodes. Equal volumes of each indexed amplicon reaction were pooled and subsequently gel-purified using the Monarch DNA Gel Extraction Kit (New England Biolabs). A single Illumina sequencing library was prepared using the xGen DNA Library Prep MC Kit (Integrated DNA Technologies) and sequenced on the MiSeq System (Illumina) at the Genome Technologies Facility (GTF) at the University of Lausanne. Data processing and analysis were performed as previously described (55).

#### Markers for fine-mapping

Insertions and deletions (INDEL) and (d)CAPS markers (**Table S8**) were designed manually based on *S. lycopersicum s2* genomic read data (23) and on *S. lycopersicum* SL4.0 (64) and *S. pimpinellifolium* LA1589 (57) reference genome sequences.

### Generation of segregating populations and recombinant inbred lines

To introduce *j2^TE^ ej2^W^* mutations in *S. pimpinellifolium* acc. LA1589, a *j2^TE^ ej2^W^* F_2_ segregant from a *s2* x LA1589 cross (23) that displayed strong suppression of inflorescence branching was selected, and their F_3_ progeny was backcrossed with LA1589 (**Fig. S1a**). One of the isolated *j2^TE^ ej2^W^* BC_1_F_2_ segregants was backcrossed with LA1589. Next, *j2^TE^ ej2^W^* BC_2_F_2_ segregants were isolated and their BC_2_F_3_ progeny was grown in the greenhouse for phenotyping.

To fine-map the *sb2* locus, *s2* x LA1589 F_2_ individuals were identified from the population used for QTL sequencing (23) that were heterozygous at the *sb2* locus based on markers chr02_39968009, chr02_41048228, chr02_43092547, and chr02_45801601, and these individuals were used as parental lines. From 768 F_3_ plants, 576 F_3_ plants with informative genotypes were identified and grown in the field at the University of Lausanne. A total of 58 recombinants that displayed either strong inflorescence branching or strong suppression of inflorescence branching were selected for the first fine-mapping population. One of the F_3_ recombinants that was homozygous at the *sb1* locus for LA1589 and segregating at the fine-mapped *sb2* locus of ∼7.4 Mb was selected as a parental line for the second fine-mapping population. For the second fine-mapping experiment, 32 F_4_ seeds were sown out from which 18 plants with informative genotypes were grown in the greenhouse. One of the F_4_ recombinants that was segregating at the fine-mapped *sb2* locus of ∼1.5 Mb was selected as a parental line for the third and fourth fine-mapping population. For the third fine-mapping population, 384 F_5_ seeds were sown out from which 112 plants with informative genotypes were selected to grow in the polytunnel at the University of Lausanne. As parental lines for the fourth fine-mapping population, the same F_4_ plant used for the third fine-mapping population was used together with a set of F_5_ plants that originated from the same F_4_ population and that had informative genotypes but for which we were not able to collect phenotyping data. A total of 240 F_5_ and F_6_ plants were sown out, from which 72 plants with informative genotypes were selected to grow in the greenhouse. For the fifth and final fine-mapping population, six F_5_ and F_6_ plants with informative genotypes were selected as parental lines. From a total of 72 F_6_ and F_7_ plants, 26 plants were identified that were homozygous at each genotyping marker within the fine-mapped *sb2* locus of ∼160 Kb and those plants were grown in the greenhouse.

To examine the effect of natural variation between domestic *s2* and wild LA1589 at the fine-mapped *sb2* locus on inflorescence branching induced by *j2^null^ ej2^null^/+* CRISPR-Cas alleles, an F_6_ individual from the fourth fine-mapping population that was homozygous at the fine-mapped *sb2* locus of ∼160 Kb for *s2* (and homozygous at the *sb1* locus for LA1589) was selected (**Fig. S5**). This *SB2* recombinant inbred line was crossed with a *j2^null^ ej2^null^*CRISPR-Cas LA1589 mutant and *j2^null^ ej2^null^ SB2* individuals were isolated from the F_2_ population. One of those selected individuals was crossed with LA1589 and from the resulting F_2_ population *j2^null^ ej2^null^/+ SB2* and *j2^null^ ej2^null^/+ sb2* individuals were grown in the greenhouse for phenotyping.

### Plant phenotyping

Inflorescence complexity was quantified by manually counting (1) the number of branching events per inflorescence, or (2) the number or branches with two or more flowers and the number of branches with less than two flowers before the first 10 branching events per inflorescence. Per replicate (individual plant), 2–7 inflorescences were counted.

### QTL sequencing analysis

To map the loci underlying suppression of *j2^TE^ ej2^W^*branching in *S. pimpinellifolium* acc. LA1589, a *s2* x LA1589 F_2_ QTL sequencing dataset obtained by (23) was re-analyzed. In (23), genomic DNA was extracted and sequenced from three pools of tissue originating from *j2^TE^ ej2^W^* double mutant F_2_ plants that displayed extreme phenotypes (A, most suppressed; B, moderately suppressed; C, branched). For this study, reads from A and B were merged into a single AB fastq file, followed by a 50% downsampling of the total number of reads using Seqtk. Reads from the AB pool, C pool, and *s2* mutant were aligned to the *S. pimpinellifolium* reference genome (LA1589_v0.1) (Glaus et al., 2025) using BWA-MEM (v.0.7.17) (65) using default parameters. Alignments were sorted and indexed using SAMtools (66) and duplicates were marked with Picard. Using the obtained BAM files and the LA1589 reference, variant calling was performed with BCFtools (66) (v.1.21; mpileup parameters: --no-BAQ --ignore-RG -d 1000000 -Q 0 --annotate FORMAT/AD,FORMAT/DP). Variants were filtered to retain high-quality, biallelic SNPs with BCFtools (v.1.21; view parameters: -i “QUAL > 50 && TYPE=’snp’ && N_ALT=1”), followed by genotype-based filtering using SnpSift (67) to retain positions homozygous for the alternate allele in the *s2* parent and homozygous for the reference allele in the LA1589 parent. The resulting VCF file was converted to tabular format using GATK, and the table was used as input for QTL sequencing analysis with the R package QTLseqR (68). Only SNPs with total read depth between 45–100 and per-pool depth of ≥ 22.5 were retained. The Δ(SNP-index) method was applied with a sliding window of 1.5 Mb, 10,000 bootstrap replications, and significance intervals of 85%, 90%, 95%, and 99% to identify genomic regions associated with phenotypic extremes. The smoothed Δ(SNP-index) values were plotted across the twelve tomato chromosomes and QTL regions were extracted.

### Transcriptome profiling

We selected two F_4_ individuals from the second fine-mapping population, one that was homozygous for *s2* and one that was homozygous for LA1589 at the fine-mapped *sb2* locus of ∼1.5 Mb. We used meristems of the F_5_ *SB2* and *sb2* recombinant inbred lines for transcriptome profiling. Meristem staging, collection, RNA extraction, and library construction were performed as previously described (19). Briefly, seedling shoot apices were collected at the transition and floral stages of meristem maturation and submerged immediately in ice-cold acetone. Shoot apices were dissected manually under a stereoscope, and three biological replicates consisting of 20–23 meristems were collected per genotype from individual seedlings. Total RNA was extracted with the Arcturus Pico-Pure RNA Isolation Kit (Thermo Scientific). Indexed libraries were prepared using the TruSeq Stranded mRNA Library Prep Kit (Illumina) and sequenced on the NovaSeq6000 System (Illumina) at the GTF at the University of Lausanne. Raw reads were aligned to the M82v1.1.0 genome sequence (Alonge et al., 2022) using STAR (v.2.7.8a) (69) and the resulting BAM files were sorted and indexed with SAMtools (66). Gene expression was quantified as unique read pairs aligned to annotated gene features (M82v.1.1.1) using HTSeq (v.2.0.3) (70). All statistical analyses of gene expression were conducted in R. Differential expression analysis was performed using DESeq2 (v.1.46.0) (71). Counts were normalized internally to account for differences in sequencing depth, and RNA composition and variance-stabilizing transformation was applied to the normalized counts. Reproducibility of biological replicates was assessed by hierarchical clustering and principal component analysis. Normalized and transformed counts were used for downstream analyses and data visualization. Differentially expressed genes (DEGs) were identified using Wald’s tests with Benjamini–Hochberg (BH) correction, considering genes with log₂ fold change ≥ 0.585 and adjusted *P* value ≤ 0.05 as significant. DEGs were clustered based on their normalized expression profiles across genotypes and meristem stages. Z-score-normalized counts were hierarchically clustered using pheatmap (v1.0.13) (72) and mean z-scores were plotted as expression trajectory across stages with cluster trends visualized using Loess-smoothing trend lines.

### Gene ontology enrichment analyses

Gene ontology enrichment analyses for biological processes were performed using a functional annotation database for the ITAG4.0 tomato genome from PLAZA (73) and the clusterProfiler package in R (74). DEG sets were tested against all annotated genes as background universe using the Benjamini-Hochberg multiple-testing correction method (pAdjustMethod = BH; pvalueCutoff = 0.1). GO term redundancy was reduced based on semantic similarity using the simplify function (cutoff = 0.7).

### Quantitative real-time PCR

First-strand complementary DNA (cDNA) was synthesized from 100 ng total RNA by qScript cDNA Synthesis Kit (Quantabio, Beverly, MA, USA). Quantitative real-time polymerase chain reaction (qPCR) reactions were carried out with a QuantStudi 6 Flex Real-Time PCR System (Thermo Scientific) using Applied Biosystems Power SYBR Green Master Mix (Thermo Scientific) and primers (**Table S8**) designed by QuantPrime (https://www.quantprime.de/) (75). Gene expression levels were quantified relative to *UBIQUITIN* (*UBI*) and *TAP42-INTERACTING PROTEIN* (*TIP41*) using the 2^-ΔΔCt^ method (Livak & Schmittgen, 2001). Statistical significance was determined by ANOVA followed by Tukey’s post-hoc analysis (*P* < 0.05).

### Protein modeling

The MADS-box tetrameric protein complexes of EJ2^WT^ and EJ2^ΔL119–S120^ were modeled using AlphaFold3. Structural visualization and geometric measurements were performed in ChimeraX. To quantify the kink angle between the two α-helices of the K-domain, an axis was defined for each helix within the tetramer. In ChimeraX, the radius of each helix was calculated as the mean atom-to-axis distance, and the angle between helix axes was subsequently computed for comparative analysis of wild-type and mutant proteins.

### Accession numbers

Sequence data from this article can be found in the EMBL/GenBank/Solgenomics data libraries under the following accession numbers: *J2* (*Solyc12g038510*), *EJ2* (*Solyc03g114840*), *STM3* (*Solyc01g092950*), *myb gene* (*Solyc02g081640*), *enhancer of polycomb* (*Solyc02g081650*), *GTP-binding* (*Solyc02g081660*), *AN* (*Solyc02g081670*), *nuclear complex protein 2* (*Solyc02g081680*), *glycosyltransferase* (*Solyc02g081690*), *proteasome subunit* (*Solyc02g081700*), *sugar transporte*r (*Solyc02g081720*), *3βHSD2* (*Solyc02g081730*), *acyltransferase* (*Solyc02g081740*), *acyltransferase* (*Solyc02g081750*), *acyltransferase* (*Solyc02g081760*), *acyltransferase* (*Solyc02g081770*), *HMGB2* (*Solyc02g081780*), *DWF4 (Solyc07g056160), SSR2 (Solyc02g069490), ROT3 (Solyc02g084740), GAME4 (Solyc12g006460), SMO4 (Solyc06g005750), C5-SD2 (Solyc02g086180), MVK (Solyc01g098840), SMO3 (Solyc01g091320), 8,7-SI (Solyc06g082980), DWF6 (Solyc10g086500), UBI (Solyc01g056940), and TIP41 (Solyc10g049850)*.

## Data availability

Raw Illumina sequence data generated in this study will be made available on SRA under the BioProject PRJNA1426717 upon publication. Seeds are available on request from S. Soyk.

## Artificial Intelligence (AI) disclosure

The authors used Microsoft Copilot and OpenAI ChatGPT solely for language editing, text polishing, and R code optimization. All outputs were verified by the corresponding author, who takes full responsibility for the analysis and conclusions.

## Acknowledgments

We thank all members of the Soyk lab, E. van der Knaap for advice on QTL mapping, and Z. Lippman for helpful discussions, providing field space and seed; J. Marquis and J. Weber from the GTF at the University of Lausanne for support with sequencing; B. Tissot, L. Nerny, A. Robadey, V. Vashanthakumar with plant care; L. Keel, A. Chatillon, T. Stupp, E. Inacio Martins, L. Héau, and S. De Mulder for support with plant care and technical support; L. Lebeigle and F. Lammers for technical support; Y. Qi for providing the pY6-SpRY vector. This work was supported by the University of Lausanne and the Fondation Herbette, the European Research Council (ERC) under the European Union’s Horizon 2020 research and innovation programme (ERC Starting Grant “EPICROP” Grant No. 802008) to S.So., and the Swiss National Science Foundation (SNSF) under an Eccellenza Professorial Fellowship (Grant No. PCEFP3_181238) to S.So.

## Author contributions

G.S. and S.So. conceived the project. G.S. and S.So. designed and planned experiments. G.S, S.A, E.La., S.St., E.Li and S.So. performed experiments and collected data. G.S., S.A, E.La. and S.So. analysed data. S.So. aquired project funding. G.S. and S.So. wrote the first draft of the manuscript. All authors read, edited, and approved the manuscript.

## Supporting information

**Table S1. Fine-mapping of *sb2* to an interval of ∼7.4 Mb.**

**Table S2. Fine-mapping of *sb2* to an interval of ∼1.5 Mb.**

**Table S3. Fine-mapping of *sb2* to an interval of ∼500 Kb.**

**Table S4. Fine-mapping of *sb2* to an interval of ∼160 Kb.**

**Table S5. Genes in the fine-mapped *sb2* interval of ∼83 Kb.**

**Table S6. Differentially expressed genes between transition meristems harvested from plants harboring the domestic *SB2* and wild *sb2* haplotype.**

**Table S7. Differentially expressed genes between floral meristems harvested from plants harboring the domestic *SB2* and wild *sb2* haplotype.**

**Table S8. Primers used in this study.**

**Table S9. gRNAs used in this study.**

**Table S10. CRISPR-Cas mutant alleles used in this study.**

## Supporting Figure Legends

**Fig. S1.**
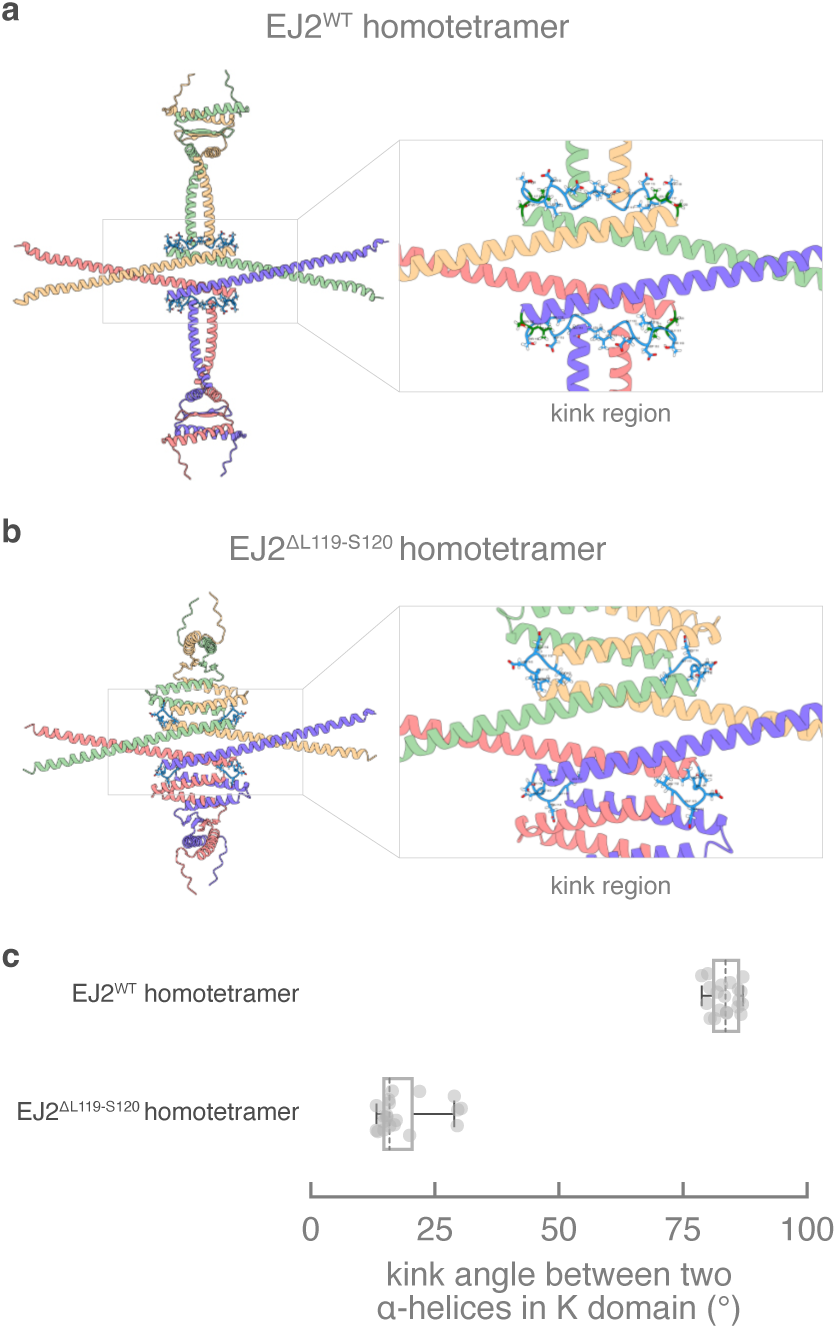
Predicted effect of the *ej2^hypo2^* allele in wild tomato on EJ2 tetramer formation. **a**,**b**, AlphaFold3-predicted MADS-box homotetramer models of EJ2^WT^ and the EJ2^ΔL119–S120^ mutant encoded by the *ej2^hypo2^* allele in wild tomato (*S. pimpinellifolium* acc. LA1589). The EJ2^ΔL119–S120^ mutant lacks two amino acid residues (L119 and S120) in the kink region that links the two α-helices of the K-domain. Each tetramer is colored by chain. The disordered C-terminal region is not shown. Boxes on the right hold close-up views of the kink region and the side chains of residues within the kink are displayed. The two residues deleted in the EJ2^ΔL119–S120^ mutant are highlighted in dark green. **c**, Quantification of the kink angle between the two α-helices of the K-domain. For each model, 20 angles were measured (*n*=20; 4 kinks per tetramer in 5 AlphaFold3 models). Boxplots represent the distribution of kink angles for EJ2^WT^ and EJ2^ΔL119–S120^. Individual EJ2^WT^ (dots) and EJ2^ΔL119–S120^ (dots) values are shown.

**Fig. S2.**
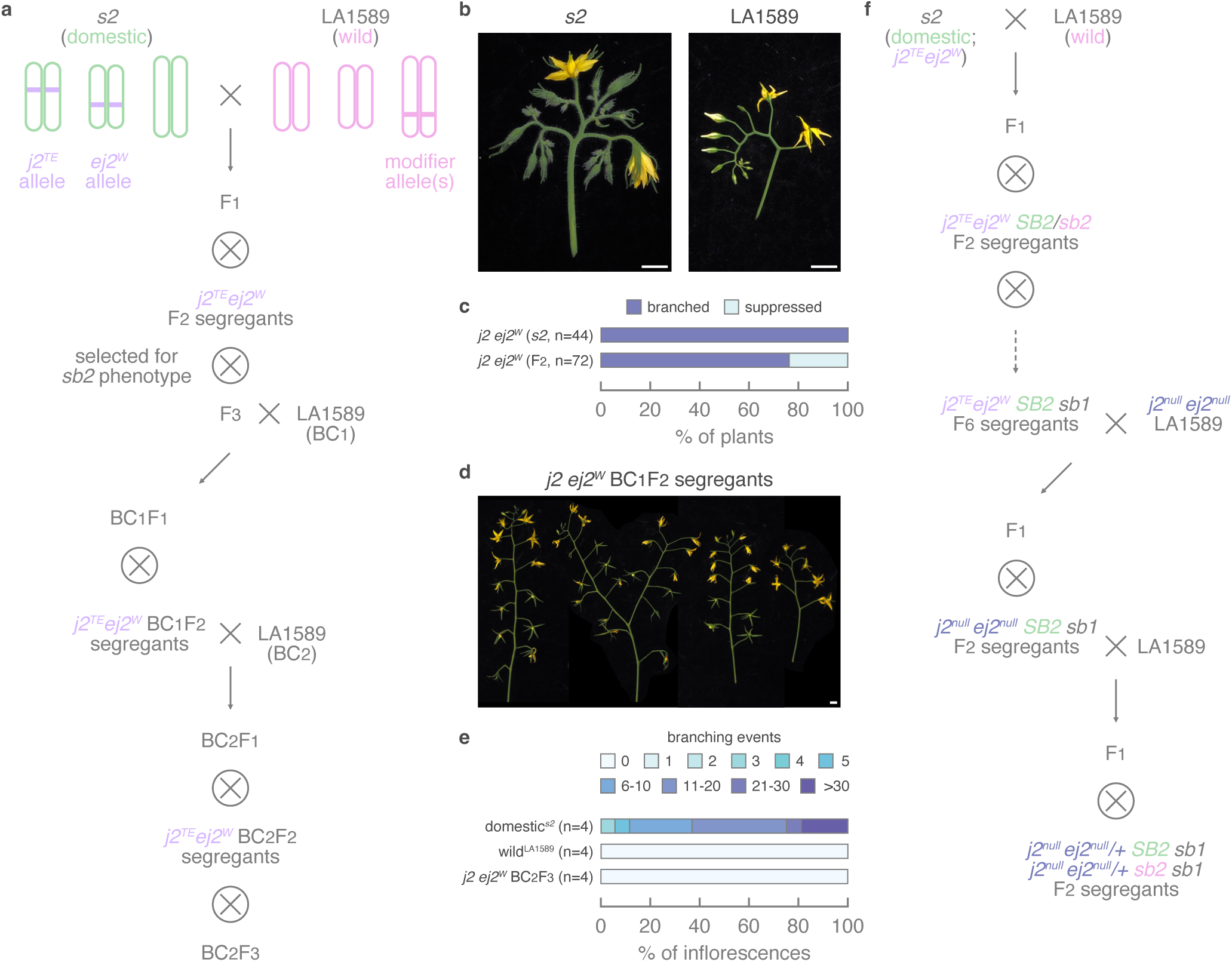
Cryptic modifier(s) in wild tomato suppress *j2^TE^ ej2^W^*-induced inflorescence branching. **a**, Crossing scheme illustrating the generation of populations used to investigate cryptic suppression of inflorescence branching of *j2^TE^ ej2^W^*-induced branching in wild tomato (*S. pimpinellifolium* acc. LA1589). The F₂ population derived from a cross between the domestic *s2* mutant (*S. lycopersicum* harboring *j2^TE^ ej2^W^*) and wild LA1589 was used for quantitative trait locus (QTL) sequencing in **Fig. S3** and Fig. 2a**,b** that identified *suppressor of branching 2* (*sb2*). To further introgress the cryptic *sb2* modifier allele(s), *j2^TE^ ej2^W^* F_3_ individuals displaying suppressed branching were repeatedly backcrossed to wild LA1589, resulting in *j2^TE^ ej2^W^* BC₂F₃ plants. **b**, Representative images of inflorescences from domestic *s2* and wild LA1589. **c**, Frequency of branched and suppressed inflorescence phenotypes among *j2^TE^ ej2^W^* F₂ plants from the mapping population. Per genotype, the number of individual plants for which inflorescence branching was evaluated is indicated by *n*. **d**, Representative images of inflorescences from *j2^TE^ ej2^W^*BC₂F₃ segregants. **e**, Quantification of inflorescence branching in *j2^TE^ ej2^W^*BC₂F₃ segregants. Per genotype, the number of individual plants for which 3-5 inflorescences were counted is indicated by *n*. **e**, Crossing scheme to generate a population that segregates both the natural *sb2* locus and the genome-edited *j2^null^*and *ej2^null^* alleles. Scalebars in **b** and **d** indicate 1 cm.

**Fig. S3.**
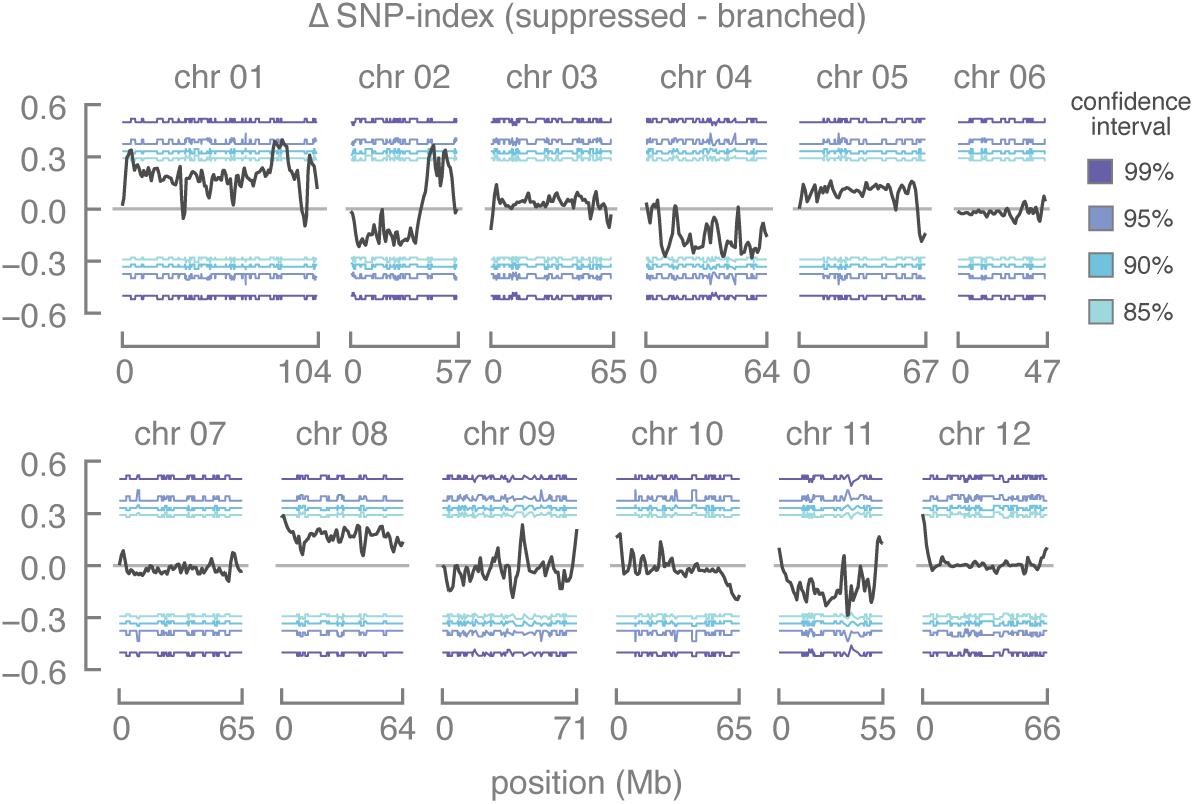
Quantitative trait loci (QTLs) for suppression of inflorescence branching in wild tomato. QTL sequencing using bulked segregants of plants with branched and suppressed inflorescences from a *s2* x LA1589 F2 population showed two *suppressor of branching* (*sb*) loci from wild tomato (*S. pimpinellifolium* acc. LA1589) on chromosomes 1 and 2.

**Fig. S4.**
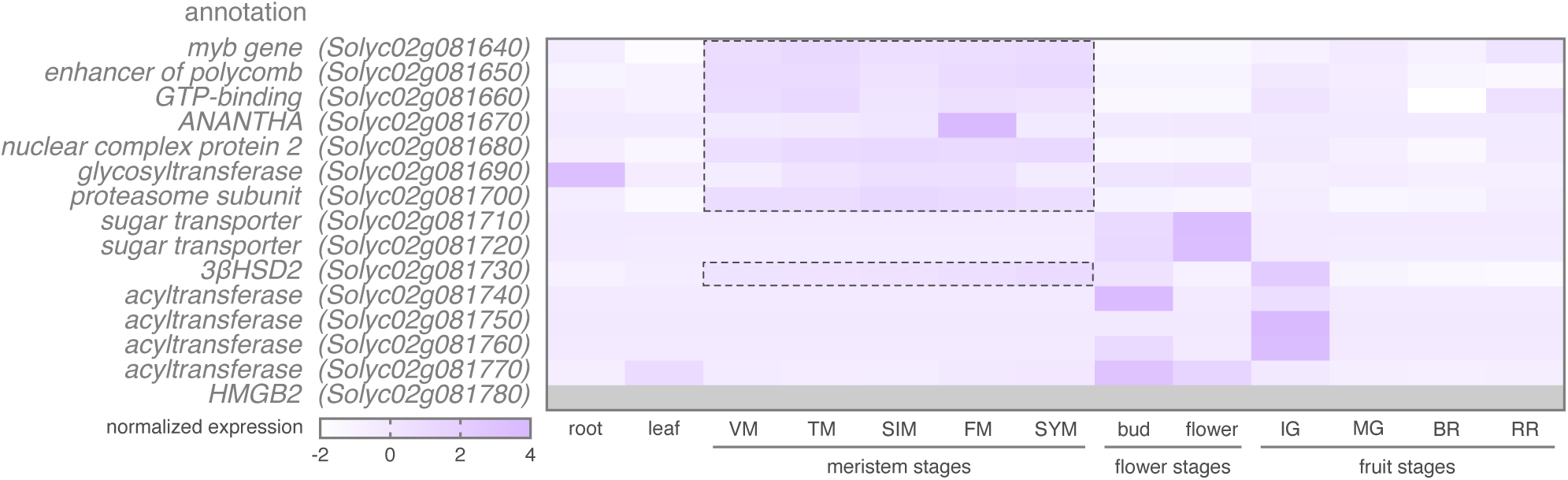
Expression profiles of genes in the fine-mapped *sb2* interval of ∼83 Kb. Normalized expression of genes in the fine-mapped *sb2* interval in different organs and developmental stages of domestic tomato (*S. lycopersicum* acc. M82). Shading represents expression relative to the highest expression value. Dotted lines mark the presence of gene expression in meristem tissues. Data were taken from the tomato meristem maturation atlas (19). Developmental stages: VM, vegetative; TM, transition; SIM, sympodial inflorescence; FM, floral; SYM, sympodial; IG, immature green; MG, mature green; BR, breaker; RR, red ripe.

**Fig. S5.**
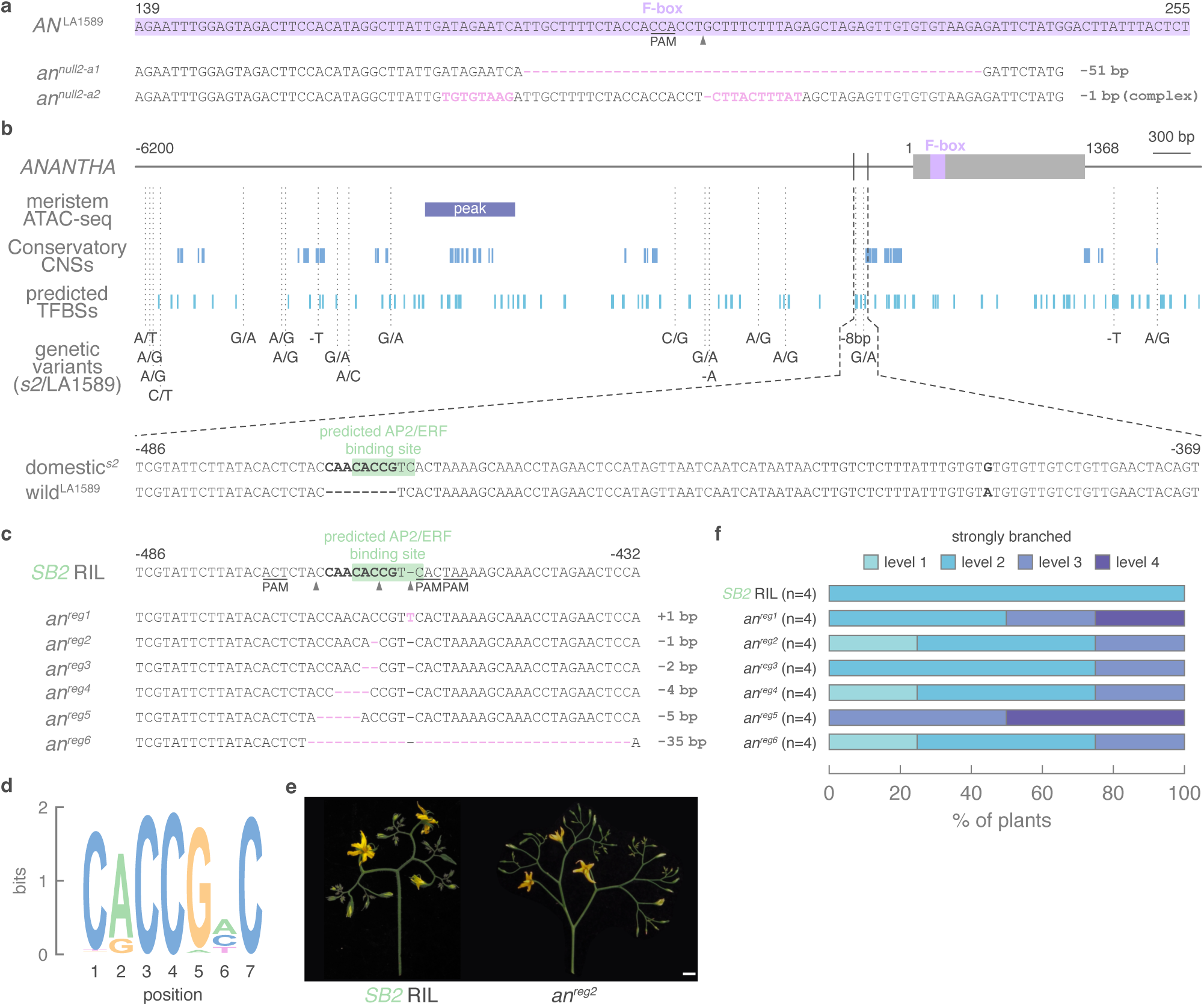
Genetic variation in coding and noncoding sequences at *ANANTHA*. **a**, Sequence encoding the F-box of the ANANTHA protein with location of the CRISPR-Cas9 target site (arrowhead) and mutant alleles in wild (*S. pimpinellifolium* acc. LA1589) tomato. **b**, A 6.2 Kb and 929 bp region upstream and downstream, respectively, of the *ANANTHA* coding region showing open chromatin, conserved non-coding sequences (CNSs), predicted transcription factor binding sites (TFBSs), and genetic variants between the natural *j2^TE^ ej2^W^* mutant (*s2*) in domestic tomato (*S. lycopersicum*) and wild tomato (*S. pimpinellifolium* acc. LA1589). A region 486–369 bp upstream of *ANANTHA* harbors a predicted AP2/ERF binding site (green) in *s2* that is disrupted by an 8-bp deletion (bold) in wild tomato acc. LA1589. **c**, CRISPR genome editing with a PAM-less Cas9 variant (SpRY) to target the predicted AP2/ERF binding site (green) in a recombinant inbred line (RIL) homozygous for the domestic *s2* mutant at the *SB2* locus. **d**, Sequence logo for the predicted AP2/ERF binding site upstream of *ANANTHA* in the domestic *s2* mutant. **e**, Representative images of inflorescences of the *SB2* RIL and *an^reg^* lines. Scalebar represents 1 cm. **f**, Semi-quantification of inflorescence branching in *an^reg^* lines. Per genotype, the number of individual plants for which the level of inflorescence branching was quantified is indicated by *n*. Sequences targeted with CRISPR-Cas in **a** and **c**: Cas9 cleavage sites for guide RNAs, PAMs, and CRISPR-Cas edits are indicated by gray arrowheads, underlining, and pink-marked text, respectively.

**Fig. S6.**
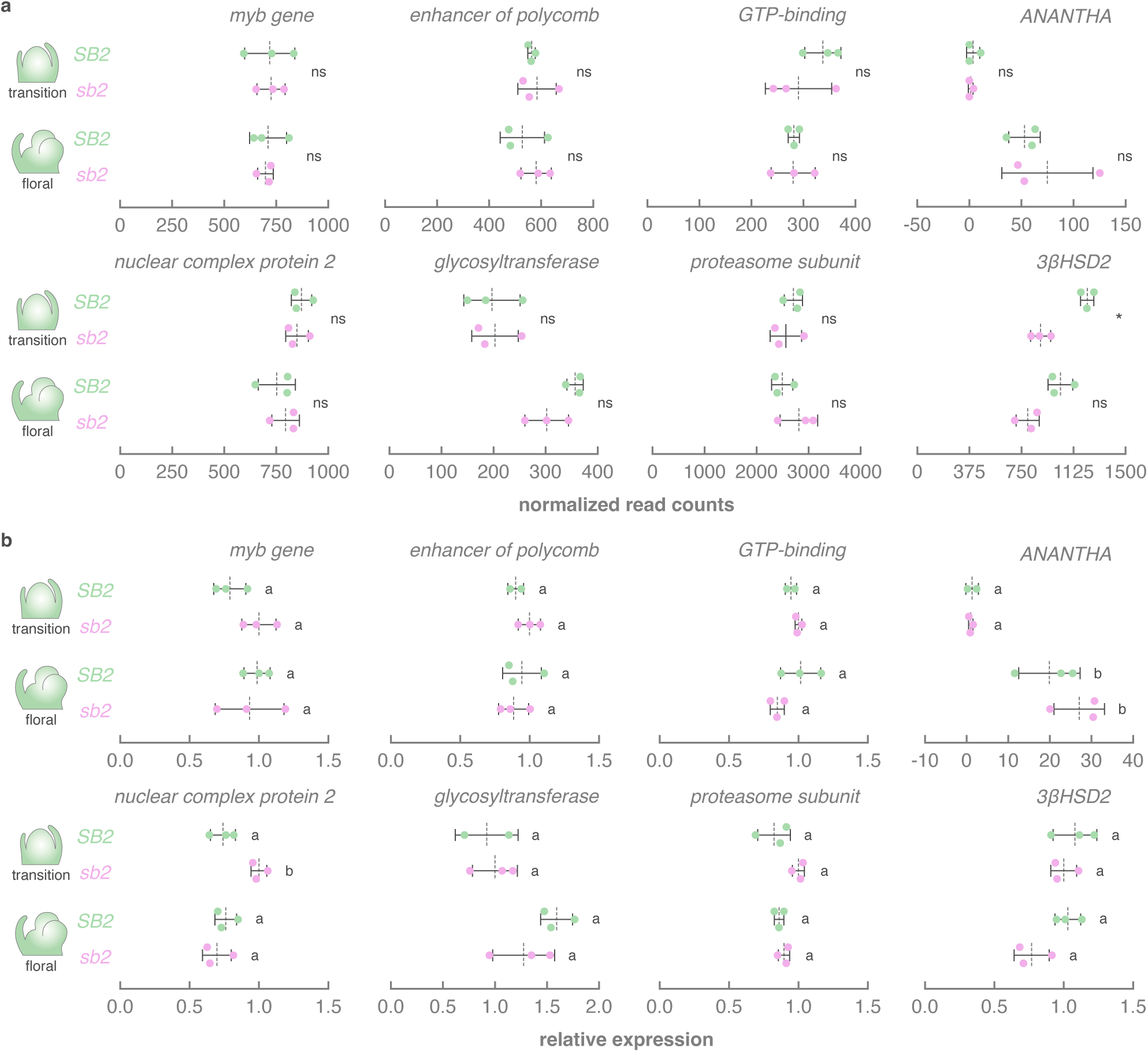
Expression analyses of *SB2* and *sb2* meristems by RNA sequencing and qPCR. **a**, Normalized expression of meristem-expressed genes in the fine-mapped *sb2* interval in transition and floral meristems from *SB2* and *sb2* plants. Dotted lines represent mean expression. Error bars denote standard deviation (*n*=3). Individual *SB2* (green dots) and *sb2* (pink dots) values are shown. Differential expression analysis was performed using DESeq2 and genes with log₂ fold change ≥ 0 and adjusted *P* value ≤ 0.05 were considered differentially expressed (indicated by *). **b**, Relative expression of meristem-expressed genes in the fine-mapped *sb2* interval in meristems at the transition and floral stage, harvested from plants harboring the domestic *SB2* or wild *sb2* locus, and analyzed by qPCR. Dotted lines represent mean expression relative to mean expression in *sb2* meristems at the transition stage. Error bars denote standard deviation (*n*=2–3). Individual *SB2* (green dots) and *sb2* (pink dots) values are shown. Statistical significance was determined by ANOVA followed by Tukey’s post-hoc analysis (*P* < 0.05; indicated by different letters).

**Fig. S7.**
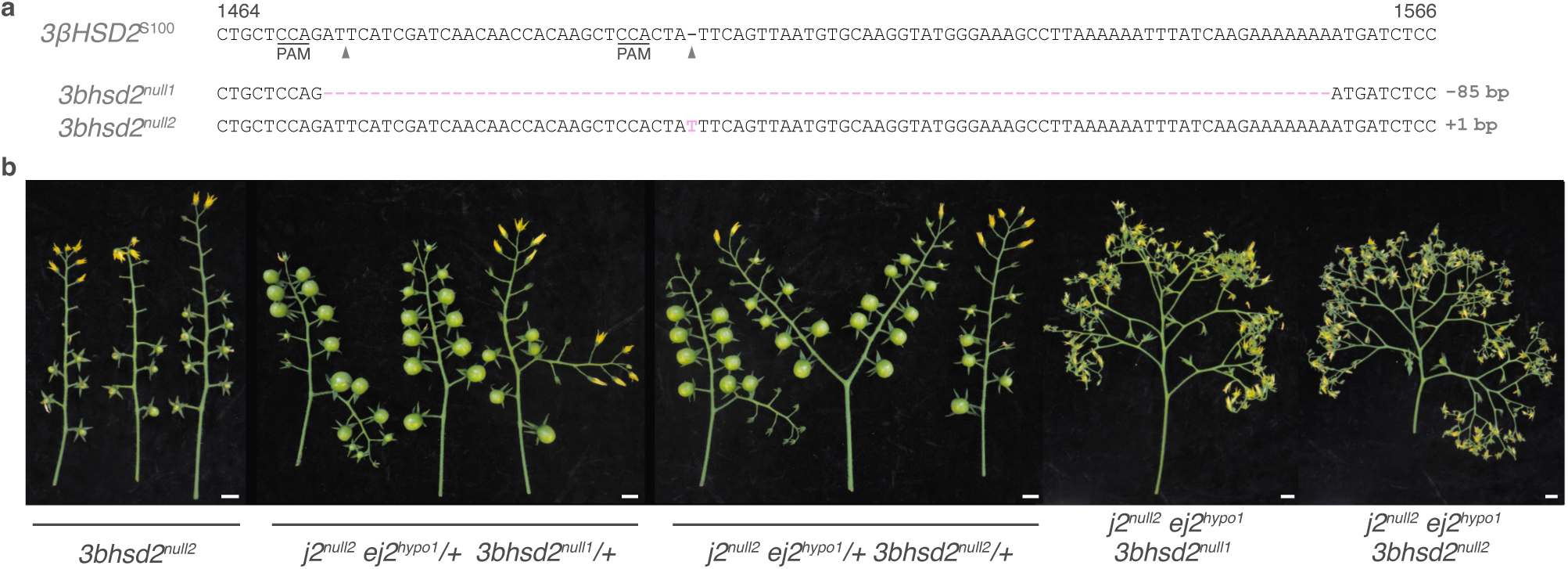
Genome editing of *3βHSD2.* **a**, Sequence of *3βHSD2* targeted by genome editing with location of the CRISPR-Cas9 target sites (arrowheads) and mutant alleles in domestic (*S. lycopersicum* acc. S100) tomato. **b**, Representative images of inflorescences from *3bhsd2^null2^*, *j2^null2^ ej2^hypo1^/+ 3bhsd2^null^/+,* and *j2^null2^ ej2^hypo1^ 3bhsd2^null^* plants. Scalebars represent 1 cm.

